# Chromosomal condensation leads to a preference for peripheral heterochromatin

**DOI:** 10.1101/714360

**Authors:** Quinn MacPherson, Andrew J. Spakowitz

## Abstract

A layer of dense heterochromatin is found at the periphery of the nucleus. Because this peripheral heterochromatin functions as a repressive phase, mechanisms that relocate genes to the periphery play an important role in regulating transcription. Using Monte-Carlo simulations, we show that an interaction between chromatin and the nuclear boundary need not be specific to heterochromatin in order to preferentially locate heterochromatin to the nuclear periphery. This observation considerably broadens the class of possible interactions that result in peripheral positioning to include boundary interactions that either weakly attract all chromatin or strongly bind to a randomly chosen small subset of loci. The key distinguishing feature of heterochromatin is its high chromatin density with respect to euchromatin. In our model this densification is caused by HP1’s preferential binding to H3K9me3 marked histone tails. We conclude that factors that are themselves unrelated to the nuclear periphery can determine which genomic regions condense to form heterochromatin and thereby control which regions are relocated to the periphery.

## INTRODUCTION

The spatial organization of chromatin in interphase eukaryotic cells is typified by a layer of dense, transcriptionally repressed heterochromatin adjacent to the nuclear periphery. This peripheral heterochromatin is visible by both electron (1) and fluorescent (2) microscopy and is present in most eukaryotes. Genes segregated to the nuclear periphery typically experience a repression of their expression (3, 4, 5, 6, 7). The highly conserved nature of peripheral heterochromatin suggests that it is a fundamental feature of nuclear architecture. Furthermore, differences in radial organization of heterochromatin could be useful as a diagnostic (8), play a role in cell differentiation (9), and—in at least the case of the rod cells in nocturnal mammals—contribute to tissue function (9).

The DNA adenine methyltransferase (Dam) technique (10, 11) identifies genomic regions that come in contact with Dam that has been attached to lamin B in human cells (12). These regions, which are typically found at the nuclear periphery (11, 12), are called Lamina Associated Domains (LADs). The position of LADs are correlated with regions of low gene density, genes with low expression levels, regions of pericentric heterochromatin, and regions with high levels of the epigenetic marks H3K9me2/3 (12, 13). However, the mechanisms that bring the LADs in contact with the nuclear periphery remain poorly understood (13, 14).

Knockouts of lamin A and lamin C—proteins that form the fibrous layer at the periphery—lead to a loss of peripheral heterochromatin (9) and a loss of peripheral positioning for tested LADs. Several additional proteins, including CEC-4, Ying-Yang 1, Emerin, Heterochromatin Protein 1 (HP1) and lamin B receptor, are shown to play a role attaching LADs to the nuclear periphery in some cell types. A review of factors implicated in position LADs is given by Steensel (13)). Cabianca *et. al*. found that cec-4 and mrg-1 independently regulate attachment of heterochromatin to the nuclear periphery in *C. elegans*, with the former providing a weak attraction and the latter anchoring heterochromatin to the periphery in an H3K9me3 independent manner (15). However, the full combination of factors that are required for positioning to the nuclear periphery remains poorly understood. For example, Ref. (9) shows the presence of lamin B receptors can cause peripheral heterochromatin in the absence of lamin A and C. Plants, which lack lamins altogether, still have peripheral heterochromatin (16). Given that LADs cover roughly a third of the human genome and peripheral heterochromatin is found in almost all eukaryotes, it is desirable to find a more general explanation for the relocation of heterochromatin regions to the nuclear periphery.

In this paper, we propose that the chromatin density difference between heterochromatin and euchromatin is capable of supplying the specificity that causes heterochromatin to localize to the nuclear periphery. Using Monte Carlo simulations, we show that a non-specific interaction with the nuclear envelope that is equally attractive of heterochromatin and euchromatin leads to peripheral heterochromatin. This envelope-chromatin interaction need only affect a small fraction of the chromatin for this effect to occur.

The higher chromatin density of heterochromatin (that is, the higher DNA and associated protein content per unit volume) is a defining feature of heterochromatin that leads to its darker appearance in stained electron microscopy.

Indeed, with 5.5 to 7.5 more DNA per unit volume (17), heterochromatin is one of the most salient features of nuclear organization. Our observations suggest that any mechanism which compacts regions of chromatin will create a preference for these regions to be segregated to the nuclear periphery. In this paper, we focus on compaction due to the preferential binding of HP1 to nucleosomes with the epigenetic mark H3K9me3 (trimethylation of the 9th lysine of the histone 3 proteins comprising the nucleosome). H3K9me3 is widely recognized as being associated with heterochromatin. Reduction in the H3K9 methylation state in the vicinity of a loci can cause its release from the nuclear periphery (18, 19, 20). Global reduction of H3K9me3 leads to loss of peripheral heterochromatin (14). While not the only source of density heterogeneity, this mark is a good starting point for understanding the heterochromatin densification and positioning in the nucleus.

To investigate the effects of this density difference on positioning of heterochromatin, we follow (21, 22, 23, 24, 25) in modeling chromatin as a block copolymer. In particular, we build on our previous work (26) where H3K9me3 levels defines the polymer blocks and densification of heterochromatin is a result of HP1-H3K9me3 binding. Our work (26) shows that this interaction leads to a phase segregation and densification of genomic regions high in H3K9me3. In this paper, we introduce into the model interactions between the chromatin beads and the nuclear periphery.

Previous simulations have included interactions between the nuclear envelope and genomic regions measured to be enriched in lamina contact to build a three dimensional model of the genome (27). Ulianov *et. al*. simulated the compaction and stretching of chromatin which comes in contact with an attractive nuclear lamin (28). Our study focus on the compaction of chromatin due to HP1-H3K9me3 binding and how the resulting compaction determines which genomic regions found at the Nuclear periphery. We demonstrate the ability of interactions which are not specific to heterochromatin to selectively reorganize the nucleus even when these interactions are relatively week or sparsely applied.

## MATERIALS AND METHODS

The phase segregation of heterochromatin and euchromatin occurs on the size scale of roughly a micron. Due to computational limitations, simulations of this scale are necessarily coarse grained. Our approach is to start with a coarse-grained model of chromatin and add a few key interactions so that consequences of each of these interactions can be investigated. Figure 1 provides a cartoon summary of the interactions included, each of which is introduced below.

**Figure 1.**
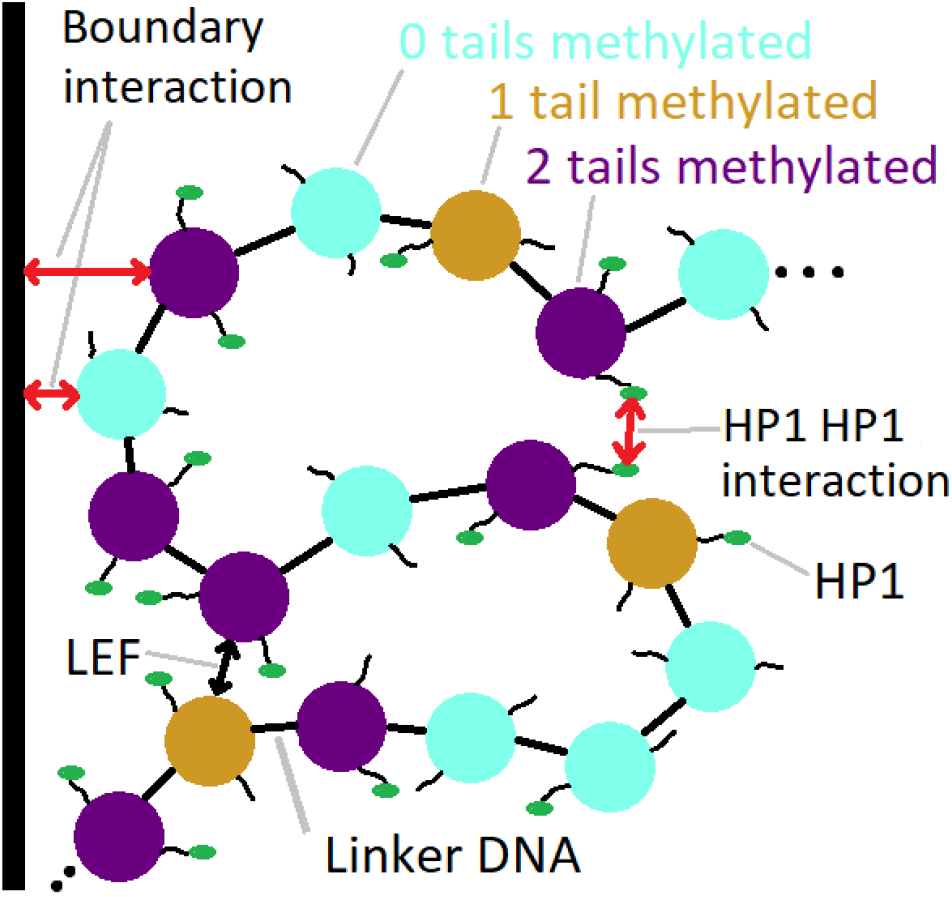
Cartoon representing the interactions included in the simulation. Nucleosomes may have neither (cyan), one (tan), or both (purple) of their tails marked with H3K9me3. Marked tails are more likely to be bond by HP1 (green oval). Interactions include HP1-HP1 binding, Loop Extrusion Factors (LEFs), and a non-specific attraction to the nuclear boundary.

### Chromatin polymer

Each computational bead in our simulation corresponds to a single nucleosome, roughly 200 bp of DNA. At this length scale, the mechanics of human chromatin is dominated by the geometry of 147 bp of DNA being wrapped around a histone octomer to form the nucleosome, separated from the following nucleosome by a roughly 50 bp long linker DNA strand. The histone octomers introduce kinks into the otherwise straight DNA backbone. These kinks, in conjunction with the natural flexibility of the linker DNA subject to thermal fluctuations, results in a Kuhn length of *ℓ_k_* = 38nm (29). The Kuhn length is defined such that for a long chromatin chain, the square of the end to end distance of the chromatin ise 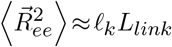 where *L_link_* is the cumulative linker length of the chromatin, i.e. *L_link_* = (16.5nm) *N_nuc_* where *N_nuc_* is the number of nucleosomes. We incorporate this result into our previous model (26) of chromatin by modeling DNA as a worm-like chain with persistence length of *ℓ_p_* = *ℓ_k_*/2=19nm with beads spaced a path length 16.5nm apart.

### Boundary interaction

For computational purposes we model a single chromosome, namely chromosome 18 of human genome assembly 19, with 390,387 nucleosome beads. Interphase chromosomes do not spread over the entire nucleus, but have been observed to segregate into separate chromosome territories (30). Therefore, we simulate chromosome 18 inside a 1.8*μ*m cube where a single face of the cube is designated as the confining nuclear periphery. This approximates the geometry of chromosome 18 within its chromosome territory interacting with the nuclear boundary as indicated by the inset of Figure 2. Chromosome 18 was chosen because it is typically found in a peripheral chromosome territory (31). Given its peripheral location, it is not surprising that chromosome 18 has a higher H3K9me3 content and lower gene content than the average chromosome.

**Figure 2.**
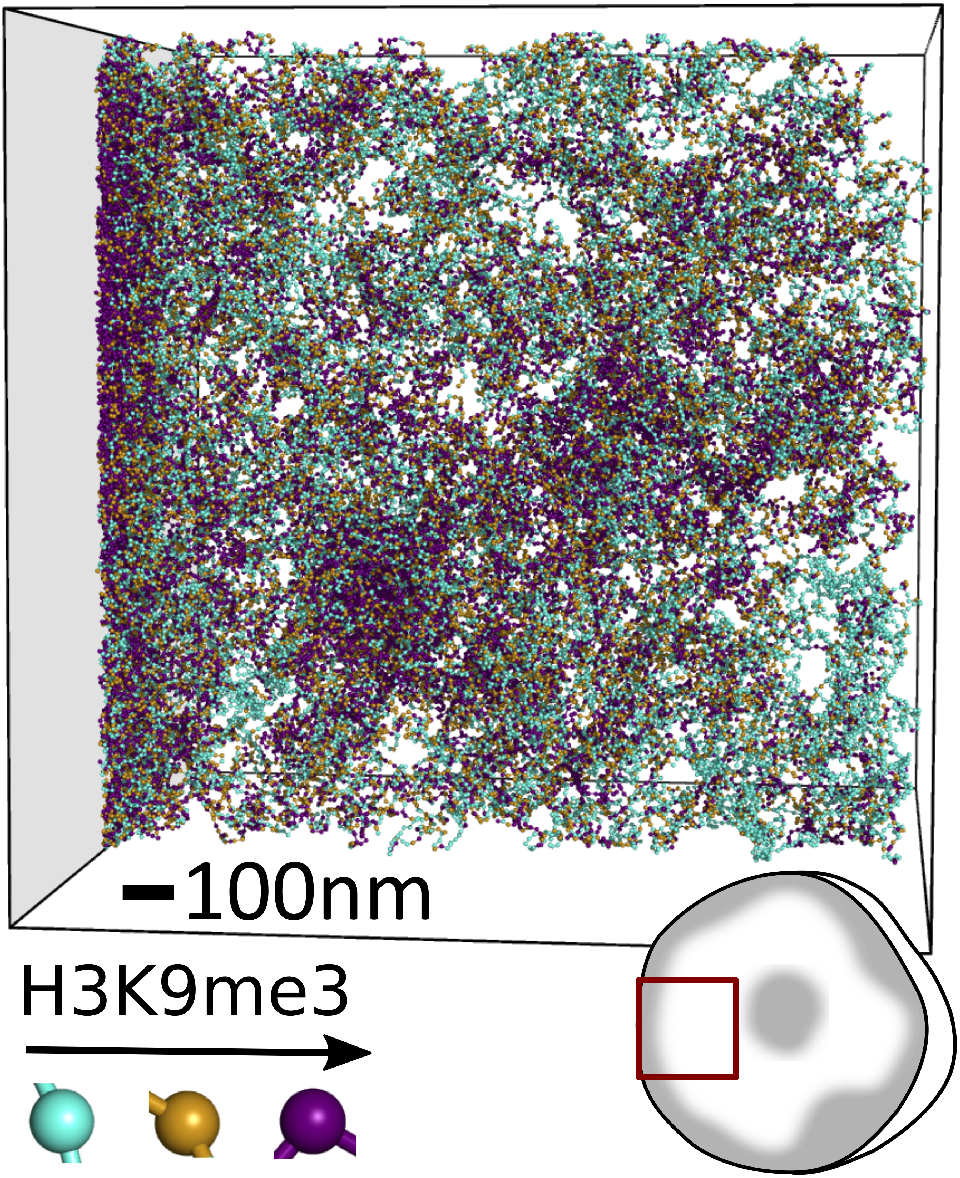
Rendering of a simulation of a single chromosome confined to a territory with non-specific attraction to the left boundary. Inset in lower left indicates approximate size in comparison to a 6*μ*m cartoon nucleus. Each bead represents a nucleosome with neither (cyan), one (tan), or both (purple) of its histone 3 tails trimethylated. Beads are only shown if they are in a 344nm (12 discretization bins) wide slice in the middle of the simulation box. Scale bar corresponds to 100nm.

### Compaction due to epigenetic modifications

Each nucleosome bead can have neither, one, or both of its histone 3 tails marked with H3K9me3. We designate the methylation state of each nucleosome based on the ChIP-seq signal for the mark (Bernstein GM12878 H3K9me3 2010, need reference) with a threshold applied such that half of all tails are marked. We choose 50% methylation because it is roughly consistent with the combined effect of the observed trimethylation and to a lesser extent dimethylation which account for 20-30% and 30-40% of the tails respectively (32, 33). The methylation pattern is held constant throughout the simulation.

The source of densification of chromatin into heterochromatin in our model is a 4.0 k_B_T oligomerization potential of HP1 coupled with a 1.53 k_B_T preference for HP1 to bind histone tails marked with H3k9me3 over those without the mark. These values are based on binding curves (34) that were analyzed in Ref. (35) to determine these energetic parameters. The effects that this condensing potential has on heterochromatin, as well as the counter balancing effects of general repulsive forces, are explored in Ref. (26). For our present purposes, it is sufficient that these interactions result in the condensation of genomic regions that are rich in H3K9me3, leaving genomic regions low in the mark to spread out to through the remaining available space.

### Loop extrusion factors

Loop Extrusion Factors (LEFs) reorganize chromatin architecture by attaching to the chromatin at two adjacent locations and then walk in opposite directions along the strand to extrude a loop of chromatin (36). The walking of the LEFs along the chromatin is inhibited in a directionally dependent manner at CTCF binding cites (36). In this way, LEFs increase the contact probability for pairs of loci which are close to each other on the genome, particularly if they are found between convergently oriented CTCF cites.

We incorporate LEFs into our simulation by requiring pairs of beads which are held together by an LEF remain spatially colocated. To determine which pairs of beads are held together by LEFs we follow Ref. (37) by running a Gillespie algorithm where LEFs stochastically walk along and fall off the chromatin and are inhibited in a directionally dependent manner by CTCF binding sites whose locations we take from Ref. (38). We assume an LEF concentration of one per 120kb and a processivity of 120kb. These numbers are chosen in accord with the best fit values found in previous studies (39). As we are using a equilibrium Monte-Carlo simulation, we first run the Gillespie algorithm to equilibrium then take the pairs of beads held together by the LEFs from the end of the Gillespie algorithm into the Monte Carlo simulation where they remain fixed. The presence of LEFs was not found to qualitatively effect our results with respect to the formation of peripheral heterochromatin (see supplemental Figure S1).

### Monte Carlo implementation

The Monte-Carlo algorithm used is an updated version of that presented in Ref. (26). The condensing HP1-based interaction, as well as a general steric repulsion, are carried out using density-based coarse-graining, where the interactions are dependent on the local density of nucleosomes *ϕ_c_*, and the local density of histone tails bound by HP1 *ρ* are each calculated using a 28.7nm discretization. The energy of oligomerization of HP1 molecules (which causes their bound nucleosomes to coalesce) is included with an energy term proportional to −*ρ*^2^. The repulsive and volume exclusion effects are included via an energy term proportional to *ϕ*^2^ as well as a hard limit *ϕ_c_* < 0.5. This density-based coarse-graining approach (40) greatly accelerates the Monte Carlo sampling used to equilibrate the structure by both reducing the computation needed to make a Monte Carlo move and softening the potentials of interaction. These effect greatly increase the probability that each move is accepted.

The same set of Monte Carlo move types from Ref. (26) where used, the most important of which is a crank-shaft move type, which rotates a section of chromatin about the axis running though its ends. In addition, a “spider” move allows semiflexible sections of DNA that are bound together by LEFs to move in a coordinate manner. Equilibration of the Monte Carlo algorithm is treated in the supplemental material of Ref. (26). Since this paper includes interactions with the boundary as well as the action of LEFs, we need to ensure that these interactions do not introduce insurmountable energy barriers that prevent the simulation from equilibrating in the roughly 10 billion crank-shaft moves used. To this end, we ensure that LEF bound nucleosomes are able to traverse the simulation confinement. To insure that the observed peripheral heterochromatin was not an artifact of the initial condition followed by poor sampling, first ran the simulation without the boundary interaction on so that heterochromatin formed away from the periphery and then turned on the boundary interaction (See supplemental Figure S2).

The FORTRAN code for our Monte-Carlo algorithm and the PYTHON code for the Gillespie algorithm are available on Github^1^.

## RESULTS

### Non-specific binding to lamina

The exact nature of the interaction(s) that bind chromatin to the nuclear periphery are unknown. That there must be at least some attraction between the two is clear. The boundary condition preventing the chromatin polymer from leaving the nuclear confinement requires the density of chromatin to approach zero outside of the confinement. Without an attraction between the chromatin and its confinement, this boundary condition, in combination with the connectivity of the DNA polymer, would lead to a region of reduced chromatin density adjacent to the boundary. In fact, in the simple model of chromatin as a long, flexible, non-interacting random walk in a spherical confinement, the density of chromatin as a function of radial distance *r* would be proportional to sin^2^ (*πr*/*r*_confine_)/*r*^2^. However, the opposite is observed, with dense heterochromatin typically being found at the nuclear boundary. This implies that there must be interaction(s) holding chromatin in proximity to the nuclear confinement.

The purpose of this paper is to demonstrate that any of a broad class of interactions will lead to peripheral heterochromatin. This potential need not be specific to heterochromatin, but can be equally attractive of all chromatin. To demonstrate this, we introduce an attractive potential of 0.3k_B_T for each nucleosome (*i.e*. independent of their methylation or HP1 binding status) when they are within the discretization bin adjacent to the nuclear lamina. The width of this potential of 28.7nm is chosen to represent the coarse-grained effect of any short range potentials, defined as potentials with a range at or below the resolution of the simulation, which is set by the discretization. If the range of the nucleosome-confinement interaction potential is much shorter, then it would need to be commensurately stronger in order to maintain the same free energy for the nucleosome to be found within the first discretization bin.

Figure 2 shows a single snap shot of the chromatin simulation with the non-specific attraction applied to the left side of the confinement box. Peripheral heterochromatin in the form of dense, H3K9me3-rich beads (purple) forms along the attractive side of the confinement. The formation of this methylation rich layer is a general feature and is not sensitive to the exact parameters chosen for the simulation, provided that a sufficiently strong non-specific attraction to the boundary is present. Also visible are regions of heterochromatin that are not in contact with the nuclear confinement, consistent with observations from microscopy. The cubical hard boundaries that prevent beads from leaving the box are meant to approximate the chromosome being confined to a single chromosome territory. Actual chromosomes are not strictly segregated into their respective territories but are able to mix to some degree. Hence, any effects of hard boundaries are artifacts of the simulation. However, they do demonstrate that hard boundaries without attractive potentials tend to be flanked by regions of less dense chromatin that is H3K9me3 depleted. This reinforces the point that some attractive potential must exist between chromatin and the nuclear confinement.

We quantify the configuration shown in Figure 2 by plotting the composition of the simulation as a function of x position in Figure 3, showing a large number of nucleosomes found within a few discretization bins of the confinement boundary. This high average density of the peripheral heterochromatin is caused by the synergistic effects of the HP1-based condensation and the attractive chromatin-boundary potential. The roughly ten-fold nucleosome density enrichment of the peripheral heterochromatin over the euchromatin is a bit higher than the 5.5-7.5 fold enrichment measured in heterochromatin (17), but this is consistent with observations of particularly dense chromatin reported at the periphery (41). This dense peripheral chromatin is notably enriched in H3K9me3, demonstrating that a non-specific binding can specifically enrich the periphery in the mark.

**Figure 3.**
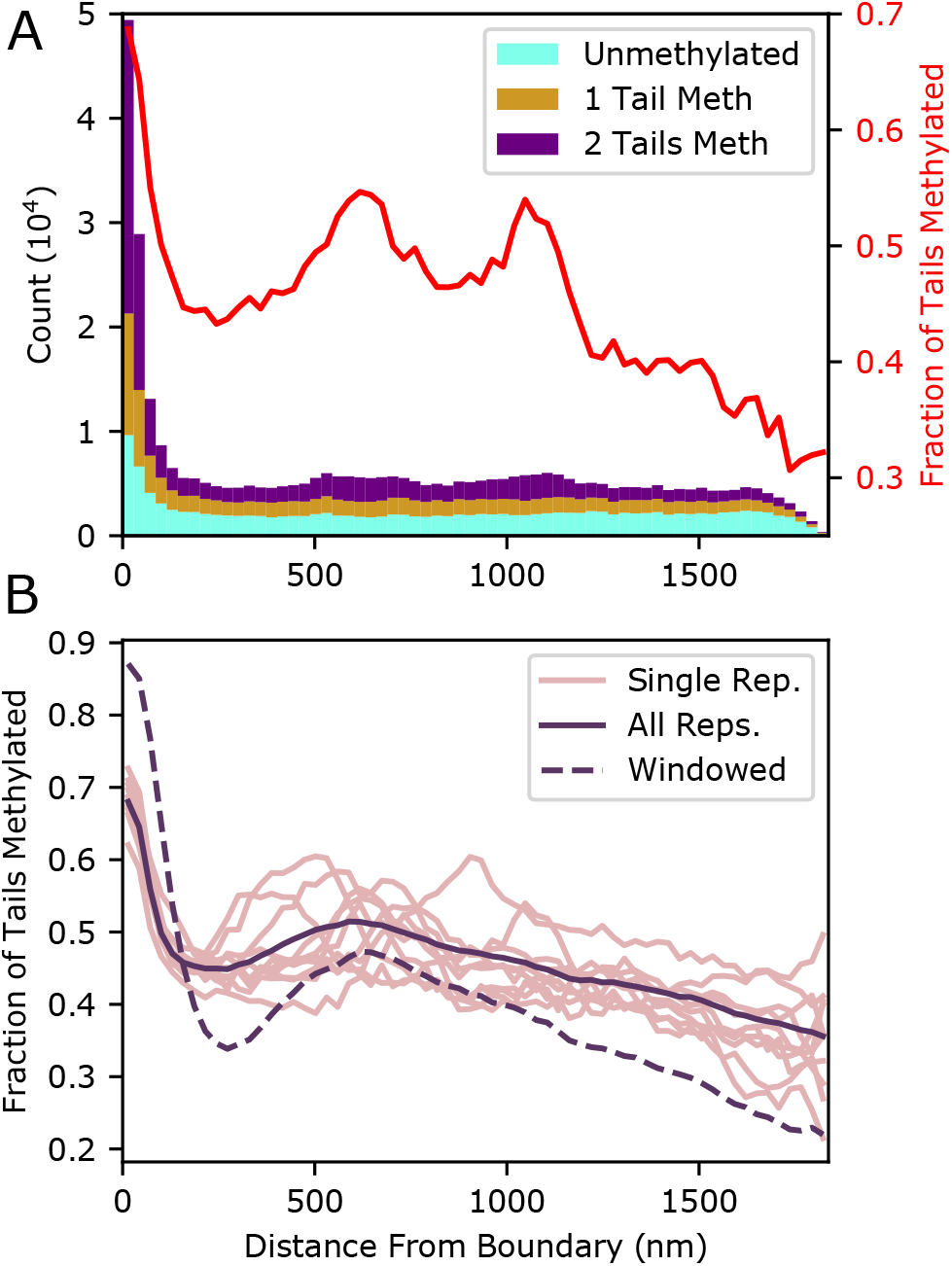
**Upper:** A stacked histogram showing the number of nucleosomes (left vertical axis) as a function of distance from the attractive boundary based on the configuration shown in Figure 2. Nucleosomes are categorized depending on whether zero (cyan), one (tan), or both (purple) of their H3 tails are trimethylated at the 9th lysine. The width of each histogram bar is a single discretization length (28.7nm). The red curve indicates the fraction of tails methylated (right vertical axis). **Lower:** Light curves: fraction of tails marked with H3K9me3, calculated using 28.7nm wide bins. Bold curve: overall fraction of tails marked for 50 configurations (10 replicates, 5 Monte Carlo configurations each). Dashed curve: fraction of tails within an H3K9me3 rich genomic region. A bead is defined as being in an H3K9me3 rich region if > 50% of the tails in the surrounding 101 nucleosome wide window (50 in either direction) have the mark.

Also visible in Figure 3 are dense, H3K9me3 rich heterochromatic regions in the interior of the simulation. The existence of non-peripheral heterochromatin is consistent with microscopy (1, 42). Note that these regions, which are also visible in Figure 2, are not quantitatively represented by the concentrations in Figure 3 that are calculated over the entire plane parallel to the boundary, only part of which contains heterochromatin.

The structure of chromatin is stochastic and continuously changing with considerable cell-to-cell variability. The configuration presented in Figures 2 and 3 are a single snap shot of a single simulation. To capture the variability inherent in this structure, we ran the Monte Carlo simulation 10 times with different positions of Loop Extrusion Factors (LEFs) and different initial conditions. The fraction-methylated profile for the final configurations of each of these simulations is shown in Figure 3b by the light curves. The overall fraction of tails methylated, calculated using the data from 5 Monte Carlo configurations from each of the 10 replicates, is shown by the solid curve in Figure 3b. An increased fraction methylated adjacent to the attractive boundary in all curves indicates the presence of H3K9me3 rich peripheral heterochromatin. Regions of non-peripheral heterochromatin—similar to those observed in Figure 2 and 3—are stochastically incorporated throughout the interior of the confinement increasing the variance in the fraction methylated. These regions of nonperipheral heterochromatin avoid all confining boundaries for entropic reasons leading to a secondary peak in the overall methylation fraction near the center of the confinement.

### Window averaging

The left end of the solid curve in 3b indicates that 66% of the nucleosomes in the immediate vicinity of the boundary are trimethylated, (compared to an overall fraction of 50%). The remaining 44% unmethylated tails—in spite of their epigenetic marking—reside in peripheral heterochromatin. Many of these are unmethylated tails that lie within otherwise highly methylated genomic regions in the ChIP-seq-based methylation profile. Figure 4 shows that it is not the methylation level of nucleosomes themselves (displayed by the coloring in the top of Figure 4), but the overall methylation of the genomic region (bottom of Figure 4) that determines peripheral positioning. The latter, which was obtained by applying a window average over the H3K9me3 of the surrounding 2kb, is more indicative of each nucleosome’s behavior than is the methylation state of the nucleosome. This observation is consistent with our early observations (26).

**Figure 4.**
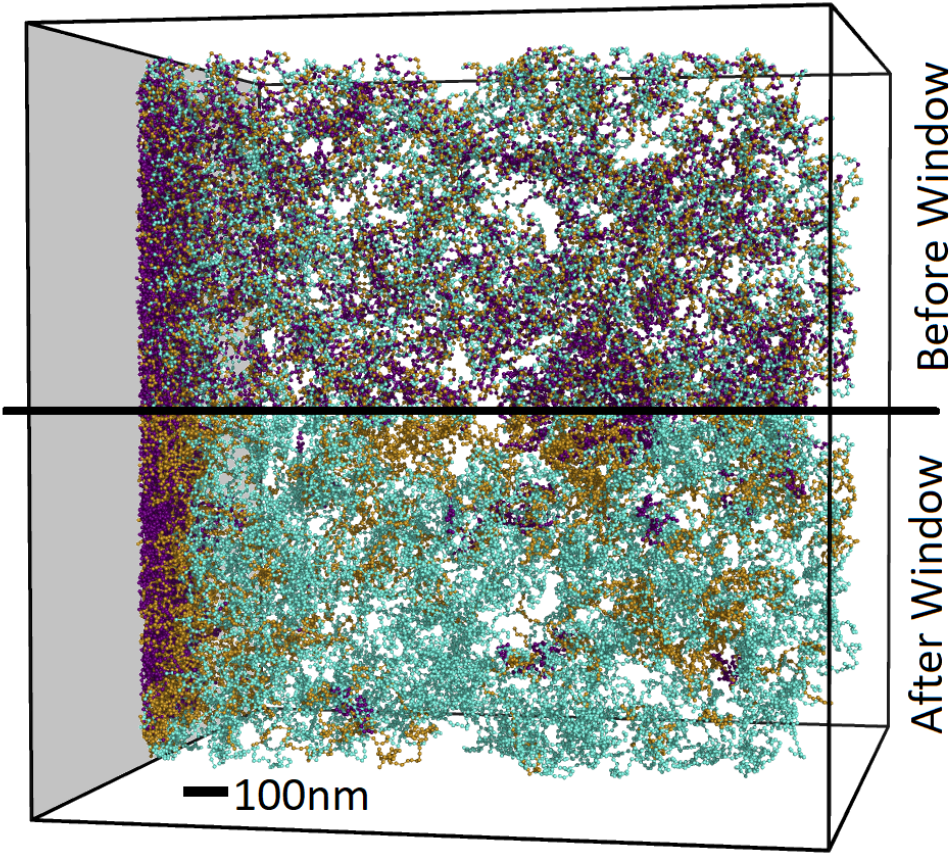
The top half of image shows our chromatin simulation with the following nucleosome coloring for the number of tails marked with H3K9me3: neither (cyan), one (tan), and both (purple). The bottom half shows the mirror image of the top half except the nucleosomes are recolored according the average methylation level of a 101-nucleosome window, with coloring indicating less than 53.0%of tails marked (cyan), 53.0-70.5%of tails marked (tan), and greater than 70.5% (purple)

To quantify the proclivity for methylation-rich genomic regions to be found at the boundary, we performed a window average of 101 nucleosomes (50 in either direction of the nucleosome of interest) of the methylation state and then reclassified the beads into H3K9me3 rich regions with more the half their tails methylated and lean regions with fewer than half their tails methylated. The fraction of nucleosomes lying at the center of methylation rich regions is shown by the dashed line in Figure 3b. Over 80% of chromatin found near the attractive boundary is considered enriched by this definition. This proportion rises to over 98% for regions with at least a third of their tails methylated. This degree of selectivity is quite significant considering that the attractive potential at the boundary is agnostic to the methylation state of the chromatin.

### Random attachments

The non-specific interaction with the nuclear confinement that is introduced in the simulation that produced Figure 2 was applied equally to all nucleosomes. While general attraction between chromatin and the nuclear confinement is sufficient to produce peripheral heterochromatin, it is not necessary. To demonstrate this we replace the chromatin-boundary potential in our simulation with a requirement that N randomly chosen nucleosomes reside within one discretization length (28.7 nm) of the left boundary. The number of beads bound N was chosen to be 75,100,150, and 300 with four replicates of each. We allow the beads to rearrange themselves on the surface by moving around within the 28.7 nm wide layer.

Figure 6 shows the fraction of tails that are methylated for the simulations with varying number of beads attached to the boundary as well as condition wide averages. As few as 150 attachments are required to cause heterochromatin to consistently form in the vicinity of the attachment boundary. Simulations with only 100 or 75 attached nucleosomes also had an increased probability of heterochromatin forming on the boundary.

We emphasize that the 150 points in Figure 5 where chosen randomly and are not correlated with the epigenetic state. These randomly selected attachments are still able to cause an H3K9me3 rich phase to form near the boundary. We observe several differences in the qualitative nature of the peripheral heterochromatin in the case of all beads being weakly attracted to the boundary (as in Figures 2, 3, and 4) versus the case of randomly attached beads (Figures 5 and Ref. 6). In the former, a layer of heterochromatin formed in the first few discretization bins adjacent to the boundary. In the latter, the droplets of heterochromatin, which are found in the interior region of the former case, are relocated to within 500 nm of the boundary. We do not intend to make a judgment about which of these cases (or perhaps a combination of them in a manner reminiscent of Ref. (15)) is most realistic *in vivo*, only to show that either is sufficient to explain the presence of peripheral heterochromatin.

**Figure 5.**
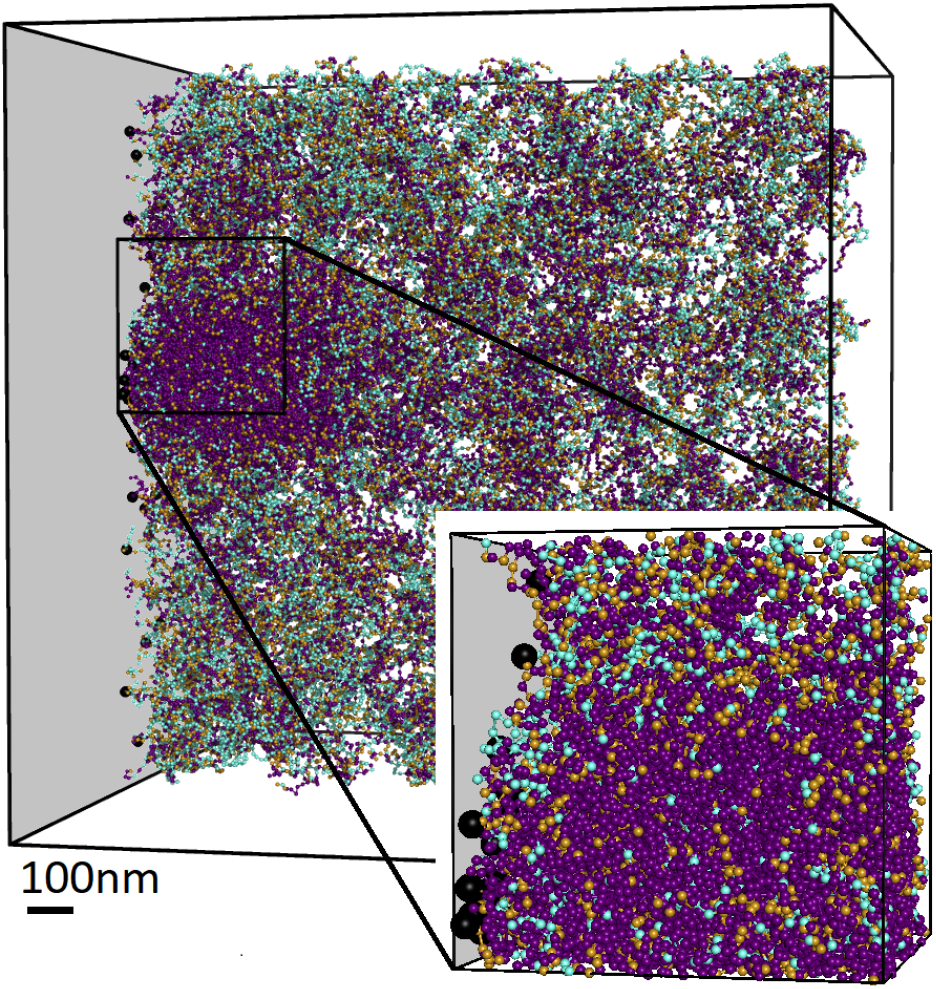
Simulation where 150 randomly chosen points (black dots) are attached to the boundary. Zoomed inset included for clarity. Note that the general attraction used in Figure 2 is no longer applied.

**Figure 6.**
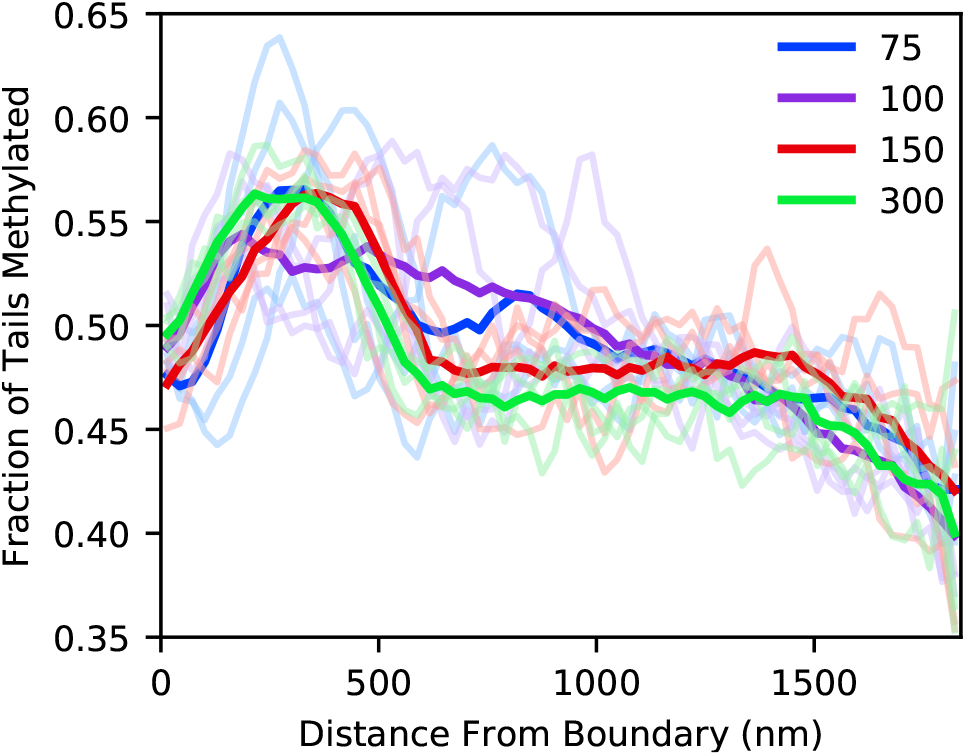
Fraction of tail methylated for configurations where a 75 to 300 beads are “bound” by requiring them to be within the first discretization. Profiles of single simulation snapshots are shown in light curves. Overall compositions for including over 4 replicates and 5 snapshots each are indicated in bold face of the same color.

### Correlation with DamID

In the model we present, H3K9me3 densifies chromatin, which can then be relocated to the nuclear periphery by nearly any attractive interaction. This observation is consistent with genomic locations high in H3K9me3 being correlated with DamID data for contact with lamin B, which is found in the fibrous layer at the nuclear periphery. Indeed, such a correlation was observed in Ref. (12, 43), providing the inspiration for the present work. Guelen’s analysis (12) focused on the H3K9me3 (and other genomic markers) in the vicinity of the boundaries of Lamin Associated Domains (LADs) which were identified as regions of enriched lamin B contact with well-defined boundaries. For our present purpose we are interested not in the boundaries of LADs *per-se*, but the overall correlation between lamin B contact and H3K9me3 measured by ChIP-seq. We therefore present a re-analysis of this correlation in Figure 7. The density of data points is indicated by the coloring of the background with darker regions corresponding to more data points. The chromosome wide averages are displayed by dots.

**Figure 7.**
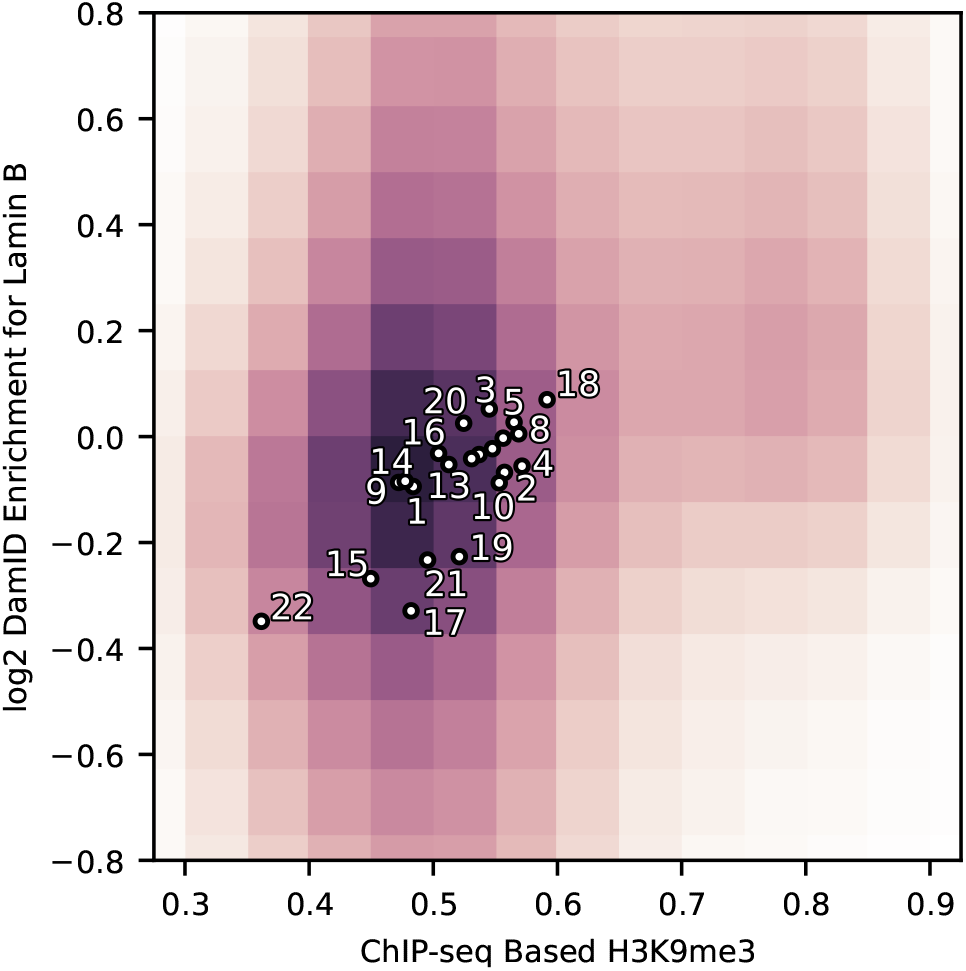
Lamin B contact is correlated with H3K9me3. The shading of the background shows the density of DamID based lamin B data points (47) and the post-cutoff ChIP-seq signal for H3K9me3 (48) of the surrounding 2kb. Dots show average DamID enrichment for corresponding chromosomes vs. the average H3K9me3 ChIP-seq signal for the entire chromosome excluding centromeres. Sex chromosomes are excluded. Unlabeled chromosomes near the center are 12, 11, 6, and 7 from left to right. The figure cropped for space purposes.

The shading in Figure 7 shows a correlation between H3K9me3 and lamin B contact. In particular there is a group of H3K9me3 rich data points (at around 0.8) with a notably enriched lamin B contact, consistent with the hypothesis of regions of H3K9me3 induced peripheral heterochromatin. Also notable is the clear correlation between chromosome wide H3K9me3 level and lamin B contact. Chromosome 18, which is the focus of our simulations, has both the highest ChIP-seq score and (apart from the sex chromosomes) the highest DamID enrichment. When observed using FISH, Chromosome 18 is typically found at the nuclear periphery (44) suggesting that H3K9me3-induced densification may also position entire chromosome territories.

*Details of lamin B DamID* vs. *H3K9me3 comparison* The DamID method (10, 11, 45, 46) identifies log2 enrichment of lamin B contact at specified loci containing the GATC motif, it is this enrichment (47) that we compare to ChIP-seq data (48) for H3K9me3. We use each of the enrichment values from all 8 data sets provided by Ref. (12) for chromosomes 1-22. The liftOver tool (49, 50) was used to convert the locations of each of these data points to hg19.

We first process the H3K9me3 ChIP-seq signal (48) by dividing it into 200 bp bins, roughly corresponding to nucleosomes. To prevent the correlation with ChIP-seq from being dominated by a small number of strongly enriched points that may be the result of biases in the ChIP-seq protocol, we apply a cutoff to the ChIP-seq signal and divide by the cutoff to put data points on a 0-1 range. This cutoff is chosen such that the post-cutoff average is one half. Based on our previous simulation work (26), we would expect the H3K9 methylation state of the surrounding roughly 2kb of chromatin to determine whether any particular GATC motif will be found in heterochromatin. We therefore perform a window average of beads 1kb (50 bins) to either side of the loci of interest. A 2D histogram of the resulting DamID - window averaged ChIP-seq data is then performed with the height of the histogram corresponding to the shading in Figure 7.

To calculate the chromosome wide averages, we averaged the post-cutoff but pre-window averaged ChIP-seq data over the entirety of each chromosome (irrespective of DamID locations), excluding centromeric regions. The average of all DamID data points is used to give the DamID value of each chromosome. We note that because the entire chromosome except the centromere is included in the average, the dot for each chromosome is not exactly the same as the average of the points that determined the shading of Figure 7. The chromosome wide average is meant to represent the chromosome’s total H3K9me3 enrichment and therefore attraction of the respective chromosome territory to the boundary.

While a clear correlation between H3K9me3 and lamin B contact is visible in Figure 7, we emphasize that the H3K9me3-HP1 interaction is far from the only mechanism which could densify chromatin and thereby cause peripheral organization. For example, the Polycomb proteins, variation in nucleosome positioning, histone tail acetylation, and supercoiling may also play a role in selectively condensing chromatin. Further muddling the correlation between H3K9me3 and lamin B contact is the difference in cell line between the two data sets, NHLF and Tig3 respectively, though both are human lung fibroblast cell lines.

## DISCUSSION

Using Monte Carlo simulations, we have shown that a nonspecific attractive force that pulls chromatin to the nuclear periphery can result in a peripheral layer of chromatin that is specifically enriched in H3K9me3. At least as important as this simulation result is the intuition it supplies for when and why peripheral heterochromatin should occur. The only difference in our simulation between nucleosomes which are marked by H3K9me3 and those that aren’t is that HP1 binding is more energetically favorable for the former. In turn, the only effect of HP1 is to oligomerize, causing tails that it is bound to (typically methylated ones) to condense into dense state (i.e. heterochromatin). For genomic regions low in H3K9me3, the entropic benefit of the freedom to move about the nucleus overpowers the condensing tendency of HP1. Without any interactions with the nuclear boundary, the heterochromatin will tend to form in the center of the confinement surrounded by euchromatin as in Ref. (26). Indeed, rod cells in the eyes of nocturnal mammals display an “inverted” organization with heterochromatin in the interior (25, 51). In that simulation this organization occurs because 1) a layer of semi-dense chromatin surrounds heterochromatin pushing it away from boundaries and 2) spreading chromatin against the confinement increases the surface area and therefore the associated surface energy of the heterochromatin phase.

If a sufficiently strong attractive force pulls chromatin to the nuclear boundary, then heterochromatin, which has a greater chromatin content per unit volume than euchromatin, will preferentially be relocated to the nuclear periphery. An alternative way to conceptualize the effect of a non-specific binding potential is to reason that an attractive boundary creates a layer of higher chromatin density. Heterochromatin, which has already given up entropy in order to condense, has less entropy to loose when it is incorporated into this boundary layer. Our Monte Carlo simulations show that a strong binding force that pulls a random subset of loci to nuclear periphery is also sufficient to explain peripheral heterochromatin. This result can be rationalized by arguing that when a small number of H3K9me3-rich nucleosomes are pulled to the boundary they pull the other beads in their phase along with them. In contrast, if a small number of H3K9me3 low loci are relocated to the boundary, they are incorporated either into or alongside the peripheral heterochromatin and don’t pull the rest of the H3K9me3 low beads—for whom they don’t have a particularly strong attraction—along with them.

Given the above arguments, we would expect a sufficiently strong non-specific attraction to cause heterochromatin to relocate to the periphery. We further argue that, given the rather low bar for such an interaction, it is not all together surprising that one (or perhaps many) exist, which is why we see peripheral heterochromatin in most eukaryotes. Indeed, we see that if as few as 150 randomly chosen loci in chromosome 18 are bound to the periphery, this is enough to cause its heterochromatin to form adjacent to the boundary. At a single such binding location for every 0.5 magabases, this is roughly an order of magnitude fewer then the average gene density of the human genome (using the rough estimate of 20,000 genes in the human genome). Alternatively, such an interaction could apply to more than 150 loci with a weaker binding affinity.

The HP1-H3K9me3 interaction is unlikely to be the only mechanism causing densification and thereby peripheral placement of chromatin. Variations in other marks such as Histone tail acetylation (52) and H3K27me (19) may also contribute. Indeed, our hypothesis predicts that any mechanism compacting chromatin (epigenetic or otherwise) should cause preferential localization to the nuclear periphery, affording a route for experimentally testing our hypothesis.

The ability to relocate heterochromatin need not apply only to the nuclear periphery. Any nuclear object to which chromatin is attracted or to which some loci are anchored could act as a condensation surface for heterochromatin. Aside from the nuclear confinement, the most prominent such object is the nucleolus. It is therefore not surprising that the nucleolus is also typically surrounded by heterochromatin. Indeed, LADs which are primarily known for their peripheral location are also associated with the nucleolus (13).

Throughout this paper, we argue that an interaction with the nuclear periphery (or nucleolus) need not be specific to heterochromatin in order to cause heterochromatin to be relocated there. While not strictly necessary, it is entirely possible that heterochromatin-specific interactions with the nuclear envelope exist in at least some organisms. Even in the presence of heterochromatin-specific interactions it is important to recognize the contribution of non-specific, density-based interactions for a number of reasons. 1) Even in a cell where a heterochromatin-specific interaction causes peripheral heterochromatin, any other mechanism which condenses a region of the genome can cause the region to be relocated to existing heterochromatin, which in turn could cause the region to be silenced. 2) Observing that some factor causes the formation of peripheral heterochromatin (or that knocking out the factor eliminates it) does not necessarily mean that the factor has any specificity for heterochromatin. 3) The search for additional mechanisms that bring heterochromatin to the periphery can be broadened to include ones without specificity for heterochromatin.

## CONCLUSION

In this paper, we investigate the effect of an attractive boundary for a system in which H3K9me3 specific HP1 binding causes the formation and densification of heterochromatin. We find that an attractive force between chromatin and the nuclear environment causes peripheral heterochromatin to form along the nuclear periphery. The interaction with chromatin need not be specific to regions high in H3K9me3 for genomic regions high in the mark to be enriched in heterochromatin. In our simulations, the formation of a dense H3K9me3 rich phase at the nuclear periphery is not sensitive to the details of the chromatin chain nor its interaction with the nuclear boundary. Peripheral heterochromatin is generated in the presence of a weak attraction between the periphery and all chromatin as well as when only a random subset of chromatin is bound to the periphery. In the latter case, heterochromatin is found to systematically form at the nuclear periphery even when less than 0.05% of all the nucleosomes are bound to the boundary, demonstrating the ability of a relatively small number of interactions to rearrange the entire nuclear structure.

We focus on heterochromatin formation based on the presence of H3K9me3. We found that the level of methylation of the 101 nucleosome long genomic region centered at a nucleosome was indicative of whether that nucleosome would be found in proximity to the nuclear boundary. Applying the 101 nucleosome window average to ChIP-seq methylation data (48) we show that it is indeed correlated with lamin B contact (12), as expected for peripheral heterochromatin. In our simulations we have assumed a fixed H3K9me3 profile based on ChIP-seq data (53). However, the spreading of the mark to neighboring nucleosomes has been observed (54), suggesting a cycle in which both H3K9me3 causes peripheral organization and the heavily H3K9 methylated environment of peripheral heterochromatin methylates nucleosomes that are relocated there. This is of great interest for future work.

Finally, we emphasize that while the interaction we introduce here is based on HP1 binding to H3K9me3 marked tails, this need not be the only source of chromatin compaction. Our results suggest that peripheral relocation can result from any interaction that densifies chromatin, epigenetic or otherwise, natural or experimentally induced.

## Supporting information

Supplemental Meterial

## ACKNOWLEDGEMENTS

We thank Bruno Beltran and Ashby Morrison for their valuable input. We thank Bradley Bernstein for publishing the ChIP-seq data and Larz Guelen for publishing the DamID data that made this work possible.

## Footnotes

1 https://github.com/SpakowitzLab/wlcsim/tree/MS2019_peripheral#chromatin-simulator

