## Supplemental Meterial for "Chromosomal condensation leads to a preference for peripheral heterochromatin"

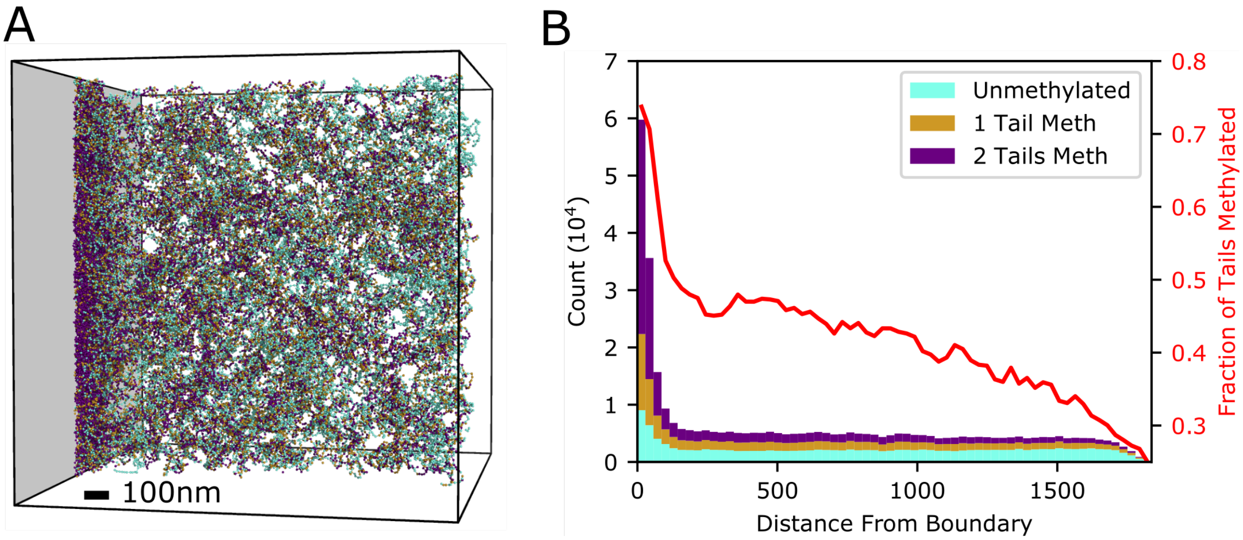


Figure S1: **A**: Simulation slice without the action of Loop Extrusion Factors (LEFs). Each bead represents a nucleosome with neither (cyan), one (tan), or both (purple) of its histone 3 tails trimethylated. **B** Composition histogram (left y axis) with colors corresponding the methylation type. Red curve (right y axis) shows fraction methylated. As in the case without LEFs, a chromatin dense, H3K9me3 rich, peripheral heterochromatin phase forms along the boundary.

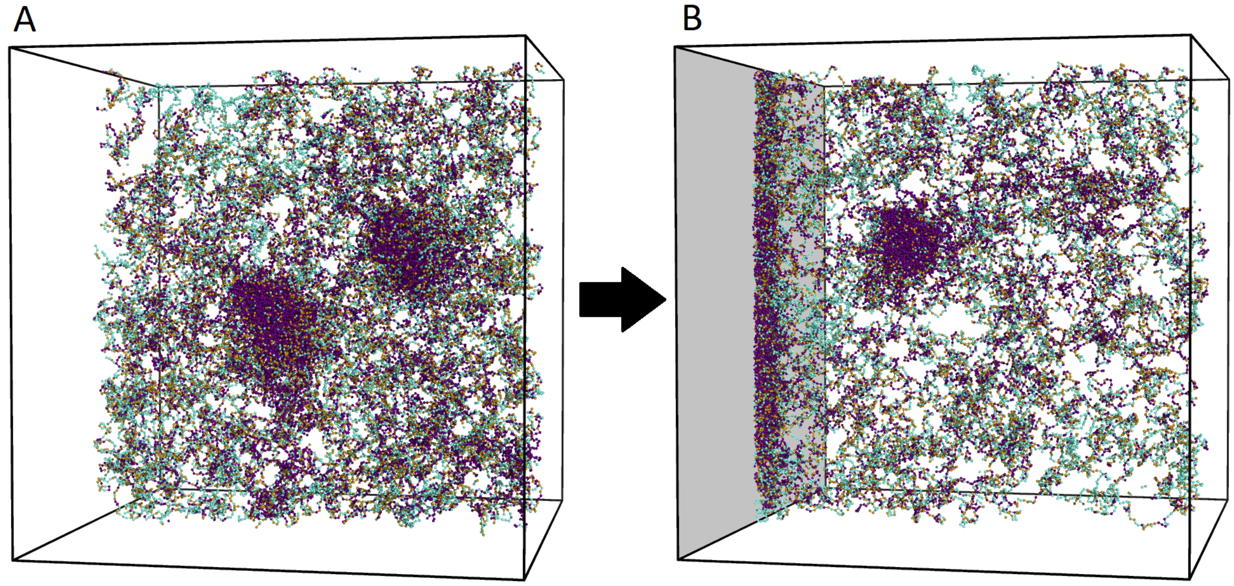


Figure S2: **A**: Simulation slice with the interaction with the boundary turned off. **B**: After turning the boundary interaction on, heterochromatin forms along the boundary. Each bead represents a nucleosome with neither (cyan), one (tan), or both (purple) of its histone 3 tails trimethylated.
